# Differential early-life survival underlies the adaptive significance of temperature-dependent sex determination in a long-lived reptile

**DOI:** 10.1101/2023.05.24.542140

**Authors:** Samantha Bock, Yeraldi Loera, Josiah Johnson, Christopher Smaga, David Lee Haskins, Tracey Tuberville, Randeep Singh, Thomas Rainwater, Philip Wilkinson, Benjamin B Parrott

**Affiliations:** University of Georgia; Princeton University; Clemson University; Tom Yawkey Wildlife Center

**Keywords:** Temperature-dependent sex determination, Reptile, Survival, Evolution

## Abstract

1. Many ectotherms rely on temperature cues experienced during development to determine offspring sex. The first descriptions of temperature-dependent sex determination (TSD) were made over 50 years ago, yet an understanding of its adaptive significance remains elusive, especially in long-lived taxa.
2. One novel hypothesis predicts that TSD should be evolutionarily favored when two criteria are met – (a) incubation temperature influences annual juvenile survival and (b) sexes mature at different ages. Under these conditions, a sex-dependent effect of incubation temperature on offspring fitness arises through differences in age at sexual maturity, with the sex that matures later benefiting disproportionately from temperatures that promote juvenile survival.
3. The American alligator (*Alligator mississippiensis*) serves as an insightful model in which to test this hypothesis, as males begin reproducing nearly a decade after females. Here, through a combination of artificial incubation experiments and mark-recapture approaches, we test the specific predictions of the survival-to-maturity hypothesis for the adaptive value of TSD by disentangling the effects of incubation temperature and sex on annual survival of alligator hatchlings across two geographically distinct sites.
4. Hatchlings incubated at male-promoting temperatures consistently exhibited higher survival compared to those incubated at female-promoting temperatures. This pattern appears independent of hatchling sex, as females produced from hormone manipulation at male-promoting temperatures exhibit similar survival to their male counterparts.
5. Additional experiments show that incubation temperature may affect early-life survival primarily by affecting the efficiency with which maternally transferred energy resources are used during development.
6. Results from this study provide the first explicit empirical support for the adaptive value of TSD in a crocodilian and point to developmental energetics as a potential unifying mechanism underlying persistent survival consequences of incubation temperature.

## Introduction

The existence of distinct sexes is a fundamental feature of nearly all metazoan taxa, however the systems by which these sexes arise are remarkably diverse (Bachtrog et al., 2014; Capel, 2017). Many taxa lack sex chromosomes and instead rely on environmental cues experienced during discrete periods in development to determine offspring sex (Bull, 1980; Devlin & Nagahama, 2002; Hobaek & Larsson, 1990). While a range of environmental factors operate in systems of environmental sex determination (ESD), including pH, nutrition, photoperiod, and social context (reviewed in (Korpelainen, 1990)), the most taxonomically widespread form of ESD is temperature-dependent sex determination (TSD) (Bull, 1980; Capel, 2017). Despite the first descriptions of TSD being made over 50 years ago (Charnier, 1966), an understanding of its adaptive significance has remained elusive.

Work by Bull and Charnov (1977) pioneered early thinking regarding the evolutionary underpinnings of TSD. Consistent with existing theory rooted in frequency-dependent selection, their model also integrated aspects of conditional sex allocation theory, with the central idea that TSD should be favored when maternal females have limited control over the environment their offspring will enter and there is a sex-dependent effect of temperature on offspring fitness (i.e., temperature-by-sex interaction) (Charnov & Bull, 1977; Shine, 1999). When these conditions are met, TSD represents an adaptive sex allocation strategy which allows females to preferentially produce the sex that will benefit most from the incubation conditions they experience. Though considerable research effort has focused on exploring sex-dependent effects of the developmental thermal environment on subsequent traits related to reproductive fitness, empirical support for such effects remains scarce. In the Atlantic silverside (*Menidia menidia*), females are produced at low temperatures characteristic of the early breeding season while males are produced at warmer temperatures characteristic of the late breeding season (Conover, 1984; Conover & Kynard, 1981). This temperature-linked difference in hatch timing leads to marked sexual size dimorphism due to females’ extended growing season. Accordingly, adult fecundity in females is more highly dependent on body size than it is in males resulting in a sex-specific effect of developmental temperature on adult fecundity (Conover, 1984). The first strong empirical evidence for the adaptive value of TSD in an amniote vertebrate came with a study in the jacky dragon, *Amphibolurus muricatus* (Warner & Shine, 2008). Using hormonal manipulations to produce males and females across a range of temperatures and quantifying lifetime reproductive success in a semi-natural field enclosure, the authors demonstrated that reproductive success of both males and females is maximized at the temperatures that normally produce each respective sex (Warner & Shine, 2008).

Demonstrating a sex-dependent effect of the developmental environment on adult fecundity in long-lived species has proven more difficult. This may indicate that TSD is adaptively neutral in these cases (Sabath et al., 2016; Valenzuela & Lance, 2004), or that adult fecundity is not the relevant target of selection (Sæther et al., 2013). Effects of the developmental environment on the other key component of fitness, survival, have received comparatively less attention. While many studies have documented temperature effects on organismal traits presumably linked to survival, including growth, morphology, and behavior (reviewed in (Noble et al., 2018)), few have quantified survival in the wild. This is especially pertinent in the context of a novel hypothesis put forth by Schwanz and colleagues which suggests TSD is evolutionarily favored when two criteria are met – (1) incubation temperature influences juvenile survival regardless of sex, and (2) sexes mature at different ages (Schwanz et al., 2016). Under these conditions, a sex-dependent effect of temperature on fitness arises through differences in age at sexual maturity, with the sex that matures later benefiting disproportionately from incubation temperatures that confer higher probability of juvenile survival. Consistent with this ‘survival-to-maturity’ hypothesis, species with TSD tend to exhibit greater sex differences in age at maturity than those in species with genetic sex determination (Bókony et al., 2019; Schwanz et al., 2016). However, a strong empirical test of this hypothesis in a field context is lacking.

The American alligator (*Alligator mississippiensis*) provides a particularly compelling system in which to assess evidence for survival-to-maturity as a potential mechanism underlying the adaptive value of TSD. Alligator embryos exhibit robust phenotypic responses to subtle changes in incubation temperature (Bock et al., 2021), and extensive nest temperature monitoring suggests maternal nesting behavior is limited in its capacity to influence incubation temperatures (Bock et al., 2020). Further, while both female and male alligators are physiologically capable of reproducing upon reaching ∼1.8 m in total length, paternity analyses demonstrate most males only begin siring offspring in the wild after reaching sizes much larger than this (e.g., > 2.8 m) (Zajdel et al., 2019), leading to a stark sex difference in age at first reproduction (∼14 years in females versus ∼24 years in males) (Wilkinson et al., 2016). Given these observations, male-promoting temperatures would be predicted to confer greater early-life survival compared to female-promoting temperatures if differential survival-to-maturity underlies the adaptive significance of TSD in this species.

The present study aimed to test the specific predictions of the survival-to-maturity hypothesis for the adaptive value of TSD in the American alligator. Toward this end, we pursued three central research questions: (1) Does incubation temperature influence early-life survival of alligator hatchlings, and specifically do male-promoting temperatures confer increased survival? (2) If so, is this due to a direct influence of incubation temperature or an effect of hatchling sex? And (3) what mechanisms mediate the lasting influence of incubation temperature on post-hatching survival? A combination of artificial incubation treatments and hormone manipulations were used to experimentally disentangle the effects of temperature from those of sex on a suite of metabolic and organismal phenotypes (**Figure 1)**. Early-life survival of hatchlings was subsequently quantified through mark-recapture approaches. Together with information about variation in alligator sex ratios across size classes throughout the species’ geographic range, results from this study provide the first explicit empirical support for the adaptive value of TSD in a crocodilian.

**Figure 1.**
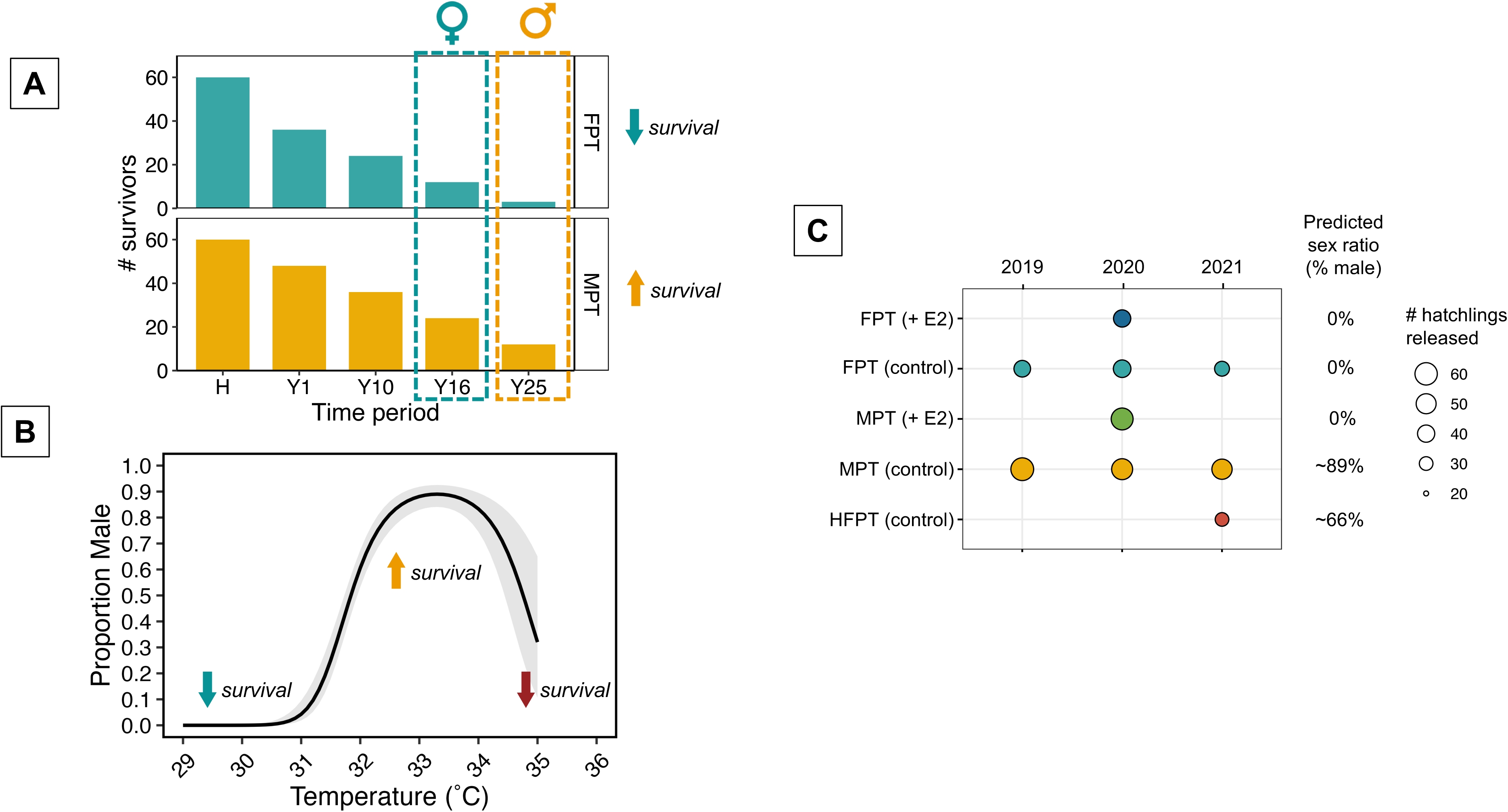
(A) Graphical summary of survival-to-maturity hypothesis for the adaptive value of temperature-dependent sex determination. (B) Temperature-by-sex ratio reaction norm of the American alligator and corresponding predictions for influences of incubation temperature on early-life survival. (C) Schematic of experimental designs implemented across study years with predicted sex ratios for each treatment group and corresponding sample sizes (number of hatchlings released for mark-recapture). Abbreviations: FPT (+E2) = female promoting temperature (29.5°C) with addition of 17ß-estradiol (0.5 μg/g egg weight); FPT (control) = female promoting temperature (29°C in 2019, 29.5°C in 2020, 2021) with either no topical treatment or addition of vehicle control (0.5 μl/g egg weight absolute ethanol; 2020 only); MPT (+E2) = male-promoting temperature (33.5°C) with addition of 17ß-estradiol (0.5 μg/g egg weight); MPT (control) = male-promoting temperature (33.5°C) with either no topical treatment or addition of vehicle control (0.5 μl/g egg weight absolute ethanol; 2020 only); HFPT (control) = high female-promoting temperature (34.5°C) with no topical treatment.

## Materials and Methods

### Summary of population sex ratios across size classes of the American alligator

Variation in American alligator population sex ratios across size classes was characterized using published data from 16 field studies of juvenile and adult size classes summarized in (Lance et al., 2000), and four studies of hatchlings and/or nest temperatures (**Table S1**). Size classes were defined based on thresholds for key life history transitions (Ferguson, 1985). Studies including a size class that spanned multiple size class definitions in the present study were excluded from analysis. Sex ratios of each size class were summarized across the species’ geographic range by taking the mean sex ratio of studies from each state (North Carolina = NC, South Carolina = SC, Louisiana = LA, Florida = FL) weighted by the reported sample size. Sexing hatchlings remains challenging in this species due to the lack of sexually dimorphic morphology in early life stages (Bock et al., 2022), thus empirically derived hatchling sex ratios were supplemented with predicted sex ratios from nest temperatures. Nest temperatures were translated to predicted hatchling sex ratios using the established temperature-by-sex ratio reaction norm for the American alligator (Lang & Andrews, 1994) and mean nest temperatures measured during the thermosensitive period, the window of time during which sex responds to temperature (Bock et al., 2020).

### Field collections and incubation experiments

All experimental procedures were approved by the Institutional Animal Care and Use Committee of the University of Georgia and field collections were permitted by the South Carolina Department of Natural Resources (SC-08-2019, SC-08-2020, and SC-08-2021). Eggs were collected from wild alligator nests across three consecutive reproductive seasons. In June 2019, nests were located via airboat at Par Pond, a 1120 ha freshwater reservoir on the Department of Energy’s Savannah River Site (PAR; Aiken, SC). Four clutches of eggs (n = 148) were collected prior to the canonical thermosensitive period of sex determination (Ferguson stage 20 – 24). In June of 2020 and 2021, nests were located via helicopter aerial surveys and subsequently accessed on foot at the Yawkey Wildlife Center (YWC; Georgetown, SC). In both 2020 and 2021, eight clutches of eggs (2020: n = 372; 2021: n = 413) were collected prior to Ferguson stage 14. All eggs were transported back to the Savannah River Ecology Laboratory (Aiken, SC) in their natal nesting material. Within 12 hours of arrival at the laboratory, a representative embryo from each clutch was staged according to (Ferguson, 1985). All eggs were candled to assess viability, weighed, and transferred to damp sphagnum moss where they were maintained at 32°C in programmable incubator chambers (Percival Scientific, model I36NLC, Perry, IA).

Experimental incubation treatments implemented in 2019, 2020, and 2021 built upon one another in a stepwise manner (**Figure 1**, **Replication statement**). To test whether hatchlings incubated at a female-promoting temperature (FPT) and male-promoting temperature (MPT) differed in their early-life survival, in 2019 eggs were distributed between two constant incubation temperatures, either 29°C (FPT; predicted 100% female) or 33.5°C (MPT; predicted ∼89% male), at Ferguson stage 17 and were maintained at these temperatures until hatching. To test whether observed differences in early-life survival between hatchlings incubated at FPT and MPT were due to incubation temperature or hatchling sex, in 2020 eggs were first distributed between two constant temperatures, either 29.5°C (FPT) or 33.5°C (MPT) at stage 15, and then received an exogenous dose of either 17ß-estradiol (E2; 0.5 μg/g egg weight; Sigma Aldrich, E2758, St. Louis, MO) or vehicle control (VEH; 0.5 μl/g egg weight absolute ethanol) at stage 19. This dose of 17ß-estradiol was chosen because it has been shown to induce complete sex reversal at MPT (Kohno et al., 2015). Thus, this 2×2 factorial design included treatment groups resulting in both presumptive males (MPT-VEH) and presumptive females from the MPT (MPT-E2), thereby disentangling incubation temperature from sex. Lastly, to test how incubation at a high-female promoting temperature (HFPT) influenced early-life survival relative to that of hatchlings from the lower incubation temperatures, in 2021 eggs were distributed at stage 15 between three constant temperatures, 29.5°C (FPT), 33.5°C (MPT), or 34.5°C (HFPT; predicted 34% female), and were maintained at these temperatures until hatching. It should be noted that temperatures above 34°C tend to produce variable sex ratios which can be male-or female-biased, however, for the purposes of this study we refer to 34.5°C as a ‘high-female promoting temperature’ to highlight its distinction from temperatures that produce consistently male-biased sex ratios. Onset (UTBI-001) HOBO temperature loggers (Bourne, MA) preprogrammed to record temperature at 5 min intervals were kept in the substrate adjacent to eggs to ensure experienced temperatures matched the intended experimental temperatures. Across all experiments, the timing at which each clutch reached key developmental stages was predicted based on the established relationship between incubation temperature and developmental rate in this species (Kohno & Guillette, 2013).

### Embryonic respirometry trials

To determine how incubation temperature influences developmental energetics in the American alligator, embryonic respirometry trials were conducted for a subset of individuals (n = 56) in 2020. Respirometry trials were conducted at stage 26, which occurs after sex determination and the approximate developmental timepoint at which metabolic rate peaks in *Crocodylus johnstoni* (Whitehead & Seymour, 1990). Trials were conducted using a flow-through respirometry system (Field Metabolic System, FMS; Sable Instruments, Las Vegas, NV) at the same temperature at which eggs were incubated, either 29.5°C or 33.5°C. Eggs were weighed just prior to the trial and then were placed in individual plastic metabolic chambers (473ml), each with an inflow and outflow channel. Metabolic chambers and an empty control chamber were kept within the FMS metabolic cooler and the constant temperature throughout the trial was controlled by a programmable PELT-5 device (Sable Instruments, Las Vegas, NV). A constant flowrate of 50 ml/min was used for all trials. All trials began between 0945 and 2200 hrs. Eggs were placed in their respective metabolic chambers and allowed to acclimate within the FMS for 1 hour prior to the initiation of measurement. Trials included one to three eggs and each egg’s metabolic chamber was measured sequentially for 25 min, with transitions between chambers controlled by an automated multiplexer (Sable Instruments, Las Vegas, NV). Consecutive metabolic measurements were separated by a 5 min measurement of the baseline control chamber and all trials ended with a 10 min baseline measurement to allow for later correction of sensor drift over the course of the trial. Data from the respirometry trials were recorded using ExpeData software (version 1.7.30; Sable Systems, Las Vegas, NV). Previously published custom scripts (Stager et al., 2021) (https://github.com/Mstager/batch_processing_Expedata_files) implemented in R statistical software version 4.1.2 were used to correct for drift in baseline O_2_ levels and extract minimum oxygen consumption (VO_2_) averaged over a 10 minute period from the raw metabolic data.

### Mark-release-recapture methods

Each individual was weighed and snout-vent length (SVL), total length, and tail girth (circumference of tail at vent) were measured upon hatching. Hatchlings were individually identified by clipping a unique pattern of keratinous tail scutes and housed together in a temperature-controlled indoor facility in custom fiberglass tanks that allowed individuals to swim and bask freely (Bock et al., 2021). Hatchlings were not fed during this period. When hatchlings reached 9–14 days old, individuals were haphazardly assigned to pods of 8–26 hatchlings each and transported back to their site of origin. Pods of hatchlings were released at a single location within ∼350 m of one another at their site of origin (PAR or YWC) over the course of three weeks. Release locations were chosen based on the availability of suitable habitat for hatchlings (e.g., presence of permanent freshwater and aquatic vegetation to provide cover) and accessibility for subsequent recapture efforts.

Monthly recapture efforts commenced at least 2 weeks after the last release date and consisted of exhaustive searches occurring on one or two consecutive nights between the hours of 1730 and 0200. Hatchlings were located visually via eyeshine (Subalusky, 2007) and were caught by hand or net from a canoe or by researchers on foot. The search area was defined as within ∼50 m of the shoreline and no further than ∼250 m from a release site. Previous studies suggest alligator hatchlings generally do not disperse more than ∼200 m during their first year of life (Deitz, 1979). Recapture efforts proceeded until hatchlings could no longer be located or captured within the search area. Recapture efforts occurred an average of four weeks apart, excluding the winter dormancy period (December – early March) during which hatchlings were assumed to be inactive (Deitz, 1979). The length of time between the last pre-winter recapture effort and first post-winter recapture effort for 2019, 2020, and 2021 was 26, 19, and 20 weeks, respectively. Monthly recapture efforts continued through the first year post-hatch.

### Data processing and statistical analyses

Statistical analyses were conducted in R version 4.1.2 (R Core Team, 2021), unless indicated otherwise. The first phase of analysis aimed at testing whether incubation temperature influences hatchling survival. Survival was assessed via two approaches. First, survival status was defined based on whether an individual was recaptured in the present time period or any subsequent time period. Survival in the pre-winter and post-winter periods were separately treated as binomial response variables and modeled using generalized linear mixed effects models (GLMMs) with a logit link function in the ‘lme4’ package (Bates et al., 2015). Candidate models explaining variation in survival included fixed effects of incubation temperature, hormone treatment (2020 only), and presumptive sex (2020 only) as well as random effects of clutch and pod identity. Coefficients of the survival GLMMs are reported as log-odds ratios (±SE), with FPT serving as the reference level in temperature comparisons for all cohorts.

Survival was also assessed by modeling individual capture histories with standard Cormack-Jolly-Seber (CJS) models in Program MARK (version 10.0) to generate maximum likelihood estimates of both apparent survival (Φ) and recapture probability (p) (White & Burnham, 1999). The use of the CJS model served to test whether observed influences of incubation temperature on survival status were due to biased recapture rates rather than apparent survival differences. Capture histories encompassed recapture efforts from October through July of the following year. Each year’s cohort was modeled separately and a weekly timestep was implemented to account for unequal sampling intervals. For all cohorts, incubation temperature was treated as a categorical grouping variable and season (pre-winter/winter, post-winter) was treated as a time-dependent covariate. For the 2020 cohort, presumptive sex was treated as an individual covariate. Models were fit using a logit-link function and candidate models were compared based on Akaike’s information criterion adjusted for small sample sizes (AICc). For the 2019 and 2021 cohorts, four candidate models were compared which variably included temperature effects on Φ and/or p. For the 2020 cohort, 16 candidate models were compared which variably included effects of temperature and/or presumptive sex on Φ and/or p. An effect of season on both Φ and/or p was included in all candidate models.

The second phase of analysis aimed at testing whether incubation temperature and/or sex reversal via exogenous estrogen treatment influences hatchling morphometric traits. The dependent variables SVL, hatchling mass, and BMI (mass/[2*SVL]) were each modeled in the ‘lme4’ package using LMMs with fixed effects of incubation temperature, hormone treatment (2020 only), presumptive sex (2020 only), and egg mass as well as a random effect of clutch identity. The third phase of analysis aimed at testing whether temperature-dependent hatchling morphometric traits explain variation in survival status. Similar to the approach described previously, survival in the pre-winter and post-winter periods were treated as binomial response variables and modeled using GLMMs with a logit link function, however in this case, candidate models included different combinations of fixed effects of hatchling morphometric traits previously shown to respond to incubation temperature. Prior to fitting these models, each trait was rescaled to have a mean of zero and standard deviation of one. Model coefficients of these LMMs are reported for the rescaled predictors. All candidate models included random effects of clutch and pod identity.

Finally, to address the extent to which incubation temperature influences the energetic cost of development, the dependent variables incubation duration (days between predicted oviposition date and pip date), embryonic metabolic rate (EMR; VO_2_), and developmental cost (product of incubation duration and EMR) were modeled with LMMs including fixed effects of incubation temperature and egg mass, and a random effect of clutch identity.

Across all analyses (apart from those implemented in program MARK), models including subsets of fixed effects were compared to the global model based on AICc using the package ‘AICcmodavg’ (Mazerolle, 2020). Models with a ΔAICc < 2.0 were considered to have support. If the best model for a response variable included a predictor with more than two levels, post-hoc comparisons between levels were conducted with the package ‘emmeans’ (Lenth et al., 2023) and p-values were adjusted according to Tukey’s HSD method. For LMMs, degrees of freedom for post-hoc comparisons were calculated according to the Kenward-Roger method. Any deviations of the global model from model assumptions (e.g., overdispersion) were diagnosed via the ‘simulateResiduals’ function in the package ‘DHARMa’ (Hartig, 2022).

## Results

### Population sex ratios become increasingly male-biased in older size classes

Population sex ratios across the geographic range of the American alligator show a consistent trend with nearly balanced or female-biased sex ratios in hatchlings and marked male biases observed in juvenile and adult size classes (**Figure 2**). In the two studies which determined the sex of over 6,000 naturally incubated hatchlings using genital morphology, yearly sex ratios ranged from 10.6 to 42.4% male (Elsey & Lang, 2014; Rhodes & Lang, 1996). Further, mean nest temperatures during the thermosensitive period in Florida and South Carolina also were predicted to yield female-biased hatchling sex ratios (40.6 and 30.6% male, respectively). In contrast, juvenile sex ratios were male-biased across each of the states for which data were available (% male, state weighted 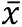 = 66.7 [NC], 60.0 [SC], 60.7 [LA], 62.6 [FL]). This was also true for adult sex ratios, with the exception of Florida (% male, state weighted 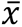 = 66.7 [NC], 61.3 [SC], 61.4 [LA], 47.2 [FL]).

**Figure 2.**
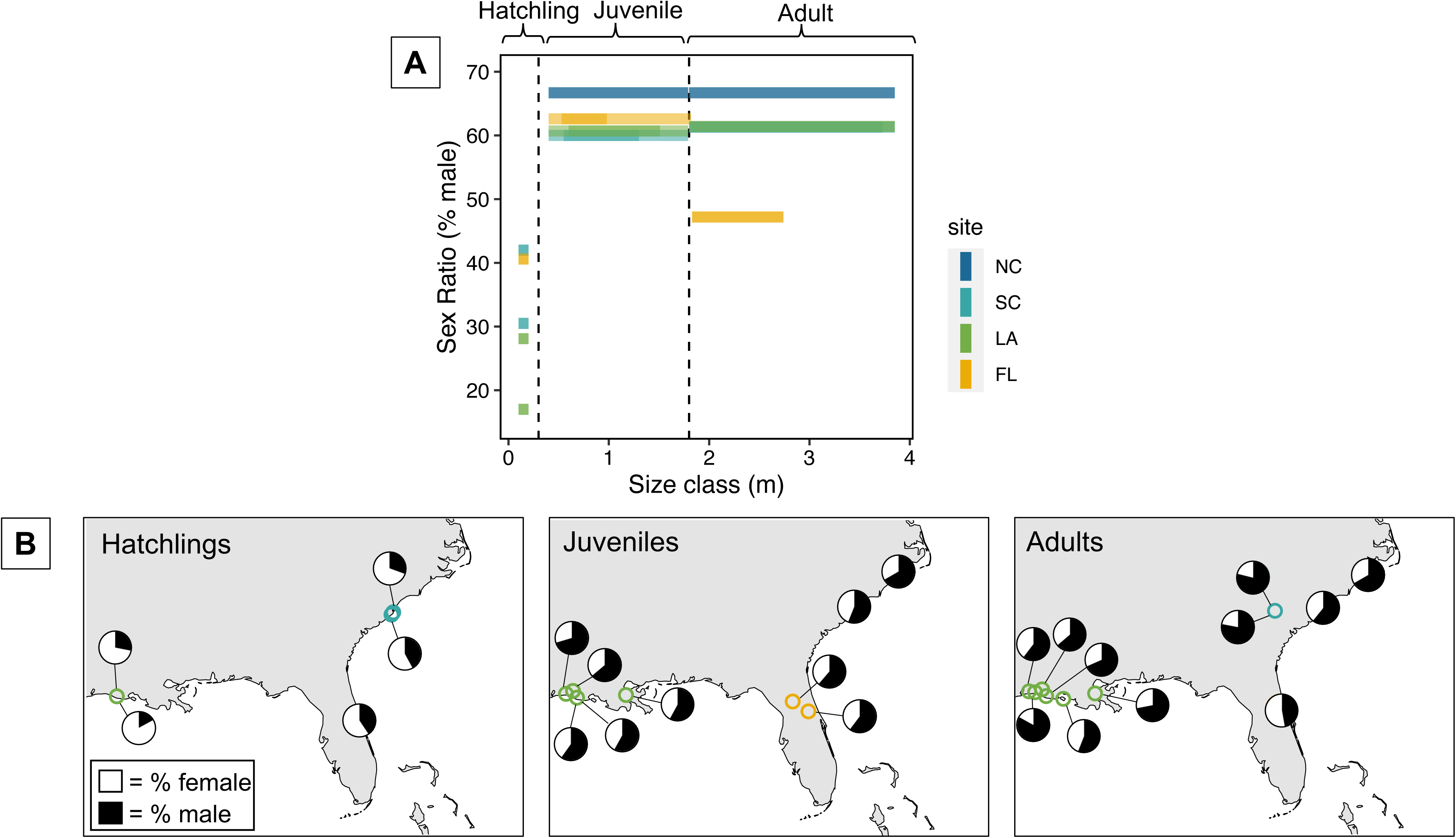
Patterns of American alligator sex ratio variation across size classes. (A) State-level weighted mean sex ratio for juvenile and adult size classes (adults defined by total length greater than 1.8m). Darker shaded regions correspond to the weighted mean maximum and minimum total length for each size class. Lighter shaded regions correspond to the absolute maximum and minimum total length included in each size class. Actual and predicted hatchling sex ratios are depicted as a single point for each individual study. (B) Sex ratios by size class across the American alligator geographic range. Each pie chart depicts a sex ratio reported by an individual study. Pie charts are associated with the locations at which each study took place. Abbreviations: NC = North Carolina, SC = South Carolina, LA = Louisiana, FL = Florida, USA.

### Hatchlings from a male-promoting temperature exhibit higher survival than those from a female-promoting temperature

Over the course of the study period, 407 hatchlings were released, and 115 unique individuals were recaptured at least once. Hatchlings incubated at MPT showed higher survival in both the pre-winter and post-winter periods compared to hatchlings incubated at FPT (**Figure 3**). In 2019, survival, as measured by recapture status, was ∼1.8 times higher for MPT hatchlings in the pre-winter period and ∼6.7 times higher in the post-winter period (**Figure 3A**). For each cohort, the GLMM that best explained variation in pre-winter survival included a fixed effect of incubation temperature (2019: ß_MPT_ = 0.91 ± 0.46; 2020: ß_MPT_ = 2.18 ± 0.78; 2021: ß_MPT_ = 1.19 ± 0.57; **Figure 3A**; **Table S2**). Though for the 2021 cohort, the null model of pre-winter survival was within 1.0 ΔAICc of the top model (ΔAICc = 0.45, *w_i_ =* 0.44). The best model explaining variation in post-winter survival for the 2019 and 2020 cohorts also included only a fixed effect of incubation temperature (2019: ß_MPT_ = 2.08 ± 1.07; 2020: ß_MPT_ = 3.48 ± 1.34; **Figure 3A**; **Table S3**).

**Figure 3.**
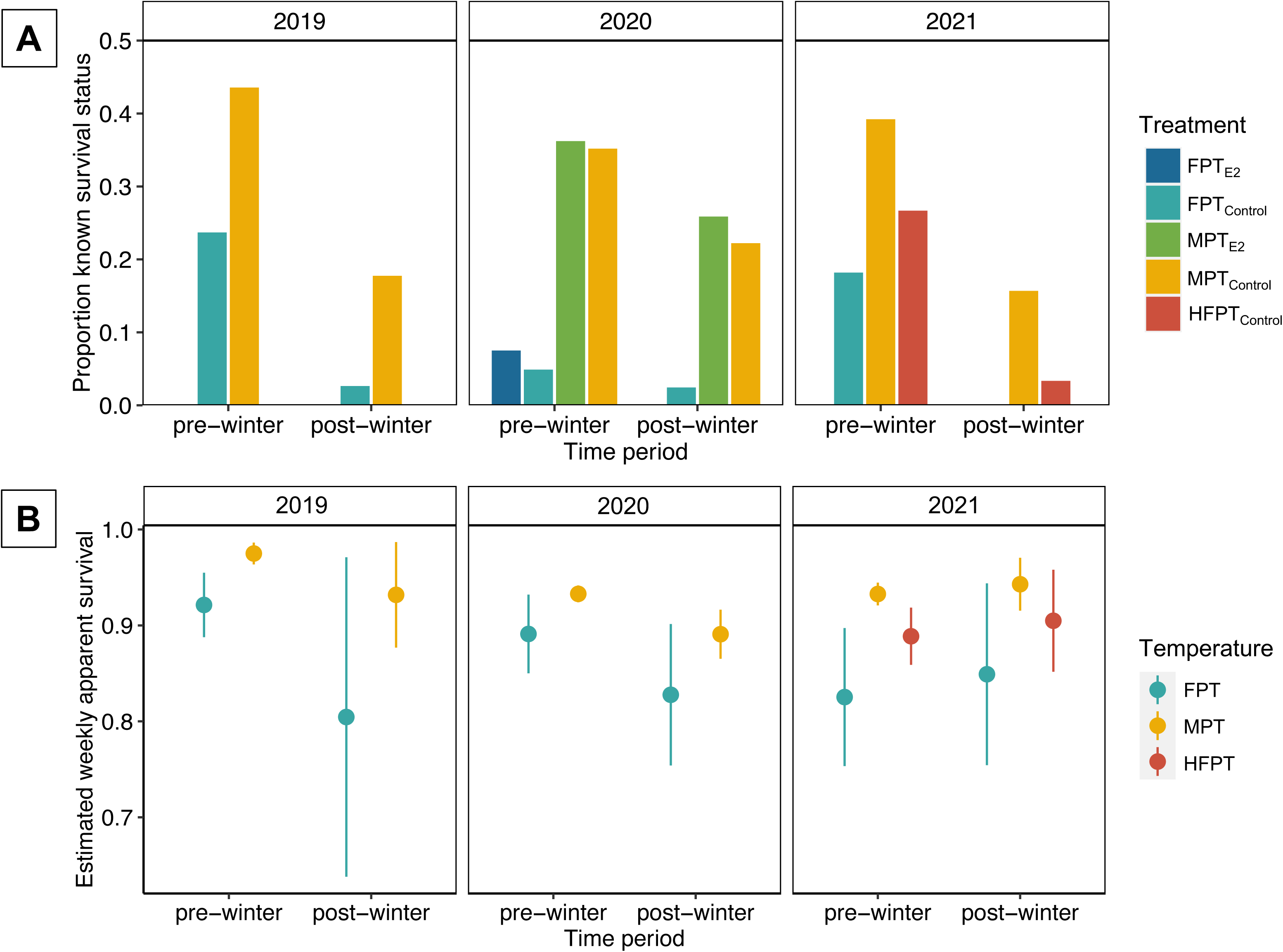
Incubation treatment effects on pre– and post-winter hatchling survival. (A) Proportion of hatchlings with known survival status in the pre-winter and post-winter periods. Known survival status was defined by an individual being recaptured at least once in the present time period or a subsequent time period. (B) Weekly apparent survival estimates from Cormack-Jolly-Seber models including effects of both temperature and season on apparent survival (Φ) and recapture probability (p). Vertical lines depict ± 1 SE. Abbreviations: FPT_E2_ = female promoting temperature (29.5°C) with addition of 17ß-estradiol (0.5 μg/g egg weight); FPT_control_ = female promoting temperature (29°C in 2019, 29.5°C in 2020, 2021) with either no topical treatment or addition of vehicle control (0.5 μl/g egg weight absolute ethanol; 2020 only); MPT_E2_ = male-promoting temperature (33.5°C) with addition of 17ß-estradiol (0.5 μg/g egg weight); MPT_control_= male-promoting temperature (33.5°C) with either no topical treatment or addition of vehicle control (0.5 μl/g egg weight absolute ethanol; 2020 only); HFPT_control_ = high female-promoting temperature (34.5°C) with no topical treatment.

In the case of the Cormack-Jolly-Seber (CJS) models, the top model for both 2019 and 2021 included an effect of incubation temperature on apparent survival, but not recapture probability (2019: ß_MPT_ = 1.23 ± 0.49; 2021: ß_MPT_ = 1.28 ± 0.32; **Figure 3B**; **Table 1**). In contrast, the top CJS model for the 2020 cohort included an effect of incubation temperature on recapture probability, but not apparent survival (ß_MPT_ = 3.77 ± 0.56; **Table 1**). Though, the model including an effect of incubation temperature on both apparent survival and recapture probability was within 1.0 ΔAICc (Φ: ß_MPT_ = 0.53 ± 0.45, p: ß_MPT_ = 3.07 ± 0.82; **Figure 3B**; **Table 1**).

**Table 1.**
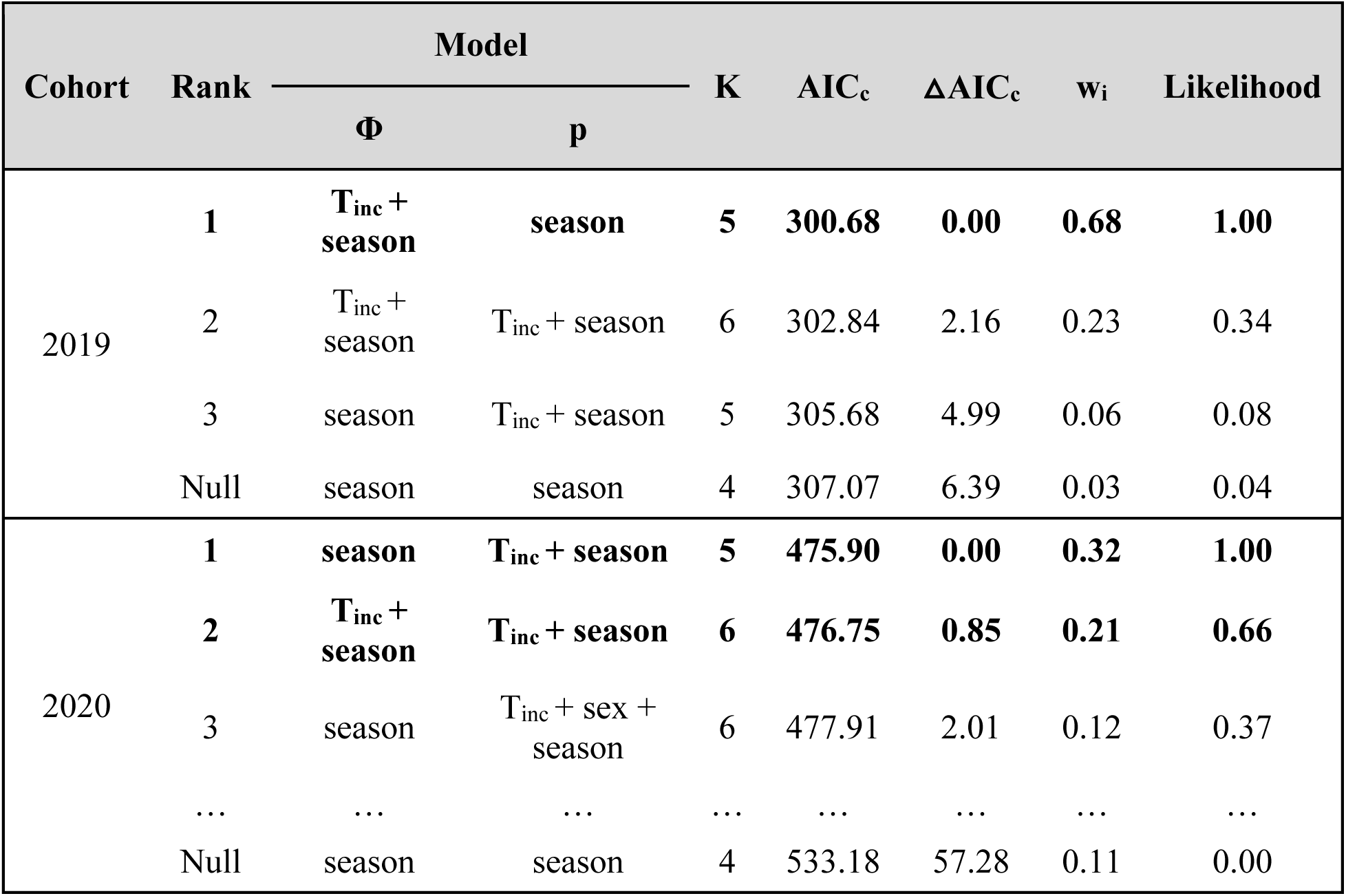
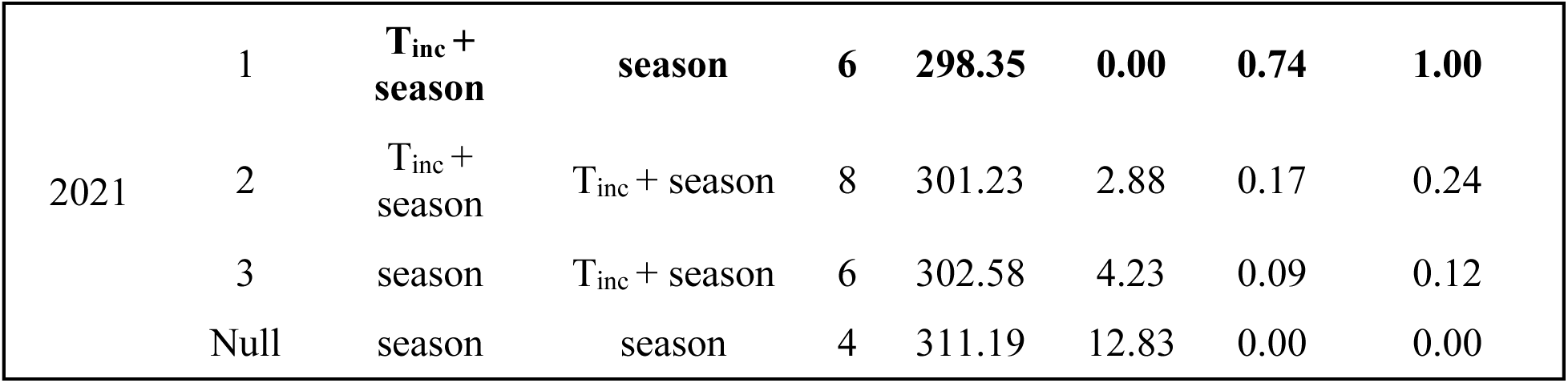
Comparison of Cormack-Jolly-Seber (CJS) models for 2019, 2020, and 2021 cohorts. All candidate models shown for 2019 and 2021 cohorts. Top three of 16 candidate models based on Akaike’s information criterion adjusted for small sample sizes (AICc) and null model shown for 2020 cohort. Abbreviations: Φ = apparent survival; p = recapture probability; K = model parameters; w_i_ = model weight; T_inc_ = incubation temperature; season = pre-winter or post-winter period.

### Incubation at MPT promotes hatchling survival independent of sex

Control male and sex-reversed female hatchlings from the MPT showed similar survival, as measured by recapture status, in both the pre-winter (Proportion surviving: MPT_Control_ = 0.35, MPT_E2_ = 0.36) and post-winter periods (Proportion surviving: MPT_Control_ = 0.22, MPT_E2_ = 0.26; **Figure 3A**). The GLMM that best explained variation in pre-winter and post-winter survival retained only a fixed effect of incubation temperature while excluding the fixed effects of presumptive sex and hormone treatment (**Figure 3A**; **Table S2,S3**). In addition, neither of the top-performing CJS models for the 2020 cohort included an effect of sex on apparent survival or recapture probability (**Table 1**). Collectively, these results suggest observed survival differences between hatchlings from the MPT and hatchlings from the FPT are due to direct effects of incubation temperature rather than a confounding influence of sex. Further, embryonic estrogen exposure does not appear to incur any consequences for hatchling survival.

### Incubation at a high female-promoting temperature confers reduced hatchling survival

Based on the shape of the alligator temperature-by-sex ratio reaction norm and the predictions of the survival-to-maturity hypothesis, high incubation temperatures that can promote the development of females (>34°C; HFPT) should confer reduced survival compared to those incubation temperatures that produce highly male-biased sex ratios (33.5°C; MPT; **Figure 1B**). Indeed, in the 2021 cohort, a lower proportion of hatchlings from the HFPT survived in the pre– winter (0.27) and post-winter (0.03) periods compared to hatchlings from the MPT (pre-winter: 0.39, post-winter: 0.16; **Figure 3A**). However, while the GLMM that best explained variation in pre-winter survival of the 2021 cohort included a fixed effect of incubation temperature, post-hoc comparisons suggest this is likely driven by the difference in survival between hatchlings from the MPT and FPT, as survival differences between hatchlings from the HFPT and MPT (log-odds ratio = –0.37, *P* = 0.78) and HFPT and FPT (log-odds ratio = 0.82, *P* = 0.45) were comparatively weaker. No hatchlings from the FPT were recaptured in the post-winter period for the 2021 cohort thereby limiting parameter estimation.

### Hatchlings incubated at MPT tend to be larger than those incubated at FPT

Incubation treatments not only contributed to variation in hatchling survival, but also to variation in hatchling morphology. Hatchling SVL was best explained by the LMM including fixed effects of both incubation temperature and egg mass for the 2019 (ß_MPT_ = 0.11 ± 0.07, ß_eggmass_ = 0.04 ± 0.01, AICc = 64.36, *w_i_ =* 0.59) and 2021 (ß_MPT_ = 0.23 ± 0.08, ß_HFPT_ = –0.69 ± 0.10, ß_eggmass_ = 0.02 ± 0.003, AICc = 112.57, *w_i_ =* 1.00) cohorts. Hatchlings from the MPT tended to have a longer SVL than hatchlings from other incubation temperatures (**Figure S1A**). Post-hoc comparisons of the three temperatures in the 2021 cohort confirmed this trend – MPT hatchlings were longer than both HFPT hatchlings (*P* < 0.0001) and FPT hatchlings (*P* = 0.03). FPT hatchlings were also longer than HFPT hatchlings (*P* < 0.0001). Incubation temperature was also associated with SVL in the 2020 cohort (ß_MPT_ = –0.30 ± 0.04), however, this effect was in the opposite direction of other years with FPT hatchlings being longer than MPT hatchlings (**Figure S1A**). Interestingly, estrogen treatment in addition to egg mass was also included in the top model for SVL (ß_E2_ = 0.19 ± 0.04, ß_eggmass_ = 0.03 ± 0.003, AICc = 79.81, *w_i_ =* 0.66), with estrogen treatment associated with longer hatchlings. SVL was the only hatchling trait for which the effect of incubation temperature was inconsistent across cohorts.

For all cohorts, the best model explaining variation in both hatchling mass and BMI included effects of incubation temperature and egg mass, with hatchlings from the MPT exhibiting consistently larger body mass (2019: ß_MPT_ = 3.78 ± 0.52, ß_eggmass_ = 0.47 ± 0.07, AICc = 481.93, *w_i_ =* 1.00; 2020: ß_MPT_ = 1.38 ± 0.35, ß_eggmass_ = 0.45 ± 0.04, AICc = 901.73, *w_i_ =* 0.54; 2021: ß_MPT_ = 4.86 ± 0.65, ß_HFPT_ = 0.59 ± 0.77, ß_eggmass_ = 0.49 ± 0.06, AICc = 585.55, *w_i_ =* 1.00) and higher BMI (2019: ß_MPT_ = 0.14 ± 0.02, ß_eggmass_ = 0.02 ± 0.003, AICc = –178.02, *w_i_ =* 1.00; 2020: ß_MPT_ = 0.11 ± 0.01, ß_eggmass_ = 0.02 ± 0.002, AICc = –326.05, *w_i_ =* 0.56; 2021: ß_MPT_ = 0.17 ± 0.02, ß_HFPT_ = 0.14 ± 0.03, ß_eggmass_ = 0.02 ± 0.002, AICc = –193.56, *w_i_ =* 1.00) compared to hatchlings from the FPT (**Figure S1B,C**). Post-hoc comparisons of the three incubation temperatures in 2021 showed that while hatchlings from the MPT were larger in mass than hatchlings from both the FPT (*P* < 0.0001) and HFPT (*P* < 0.0001), hatchlings from the FPT and HFPT did not differ from each other (*P* = 0.73). In the case of BMI, however, hatchlings from the MPT and HFPT showed higher BMI than hatchlings from the FPT (*P* < 0.0001), but BMI did not differ between the MPT and HFPT (*P* = 0.42). Presumptive sex was not included in the top model for any of the traits examined. Overall, temperature consistently exerted a strong influence on hatchling morphology, and specifically, incubation at MPT promoted the development of larger hatchlings with greater body condition (as indicated by BMI). Given this observation, further analyses were aimed at determining whether these temperature-related traits might explain variation in hatchling survival.

### Survival is weakly associated with temperature-dependent hatchling morphometric traits

If incubation temperature drives differences in survival through its persistent effects on hatchling traits, variation in temperature-related traits should correspond to variation in survival. In the 2019 and 2021 cohorts, pre-winter survival was best explained by variation in hatchling SVL, wherein longer hatchlings were more likely to survive (2019: ß_SVL_ = 0.44 ± 0.23, AICc = 135.22, *w_i_* = 0.27; 2021: ß_SVL_ = 0.56 ± 0.30, AICc = 139.87, *w_i_* = 0.32; **Figure 4A**). The second-best model of pre-winter survival in the 2019 cohort included a single effect of body mass, with larger hatchlings again showing a survival advantage over smaller hatchlings (ß_MASS_ = 0.37 ± 0.21, AICc = 136.08, *w_i_* = 0.18; **Figure 4B**). Though, for both cohorts, the null model was within 2 ΔAICc of the top models (2019: ΔAICc =1.71, *w_i_=* 0.12; 2021: ΔAICc =1.80, *w_i_=* 0.13). Further, post-winter survival was not well explained by any of the temperature-related traits examined. Results were similarly mixed for the 2020 cohort. While pre-winter survival was not well explained by any hatchling traits, the top model for post-winter survival included an effect of BMI (ß_BMI_ = 0.73 ± 0.35, AICc = 134.41, *w_i_* = 0.29), with higher condition hatchlings exhibiting greater survival (**Figure 4C**). While a few associations between survival and temperature-related hatchling morphometric traits were detected, no single trait emerged as a strong candidate for mediating the effect of incubation temperature on survival.

**Figure 4.**
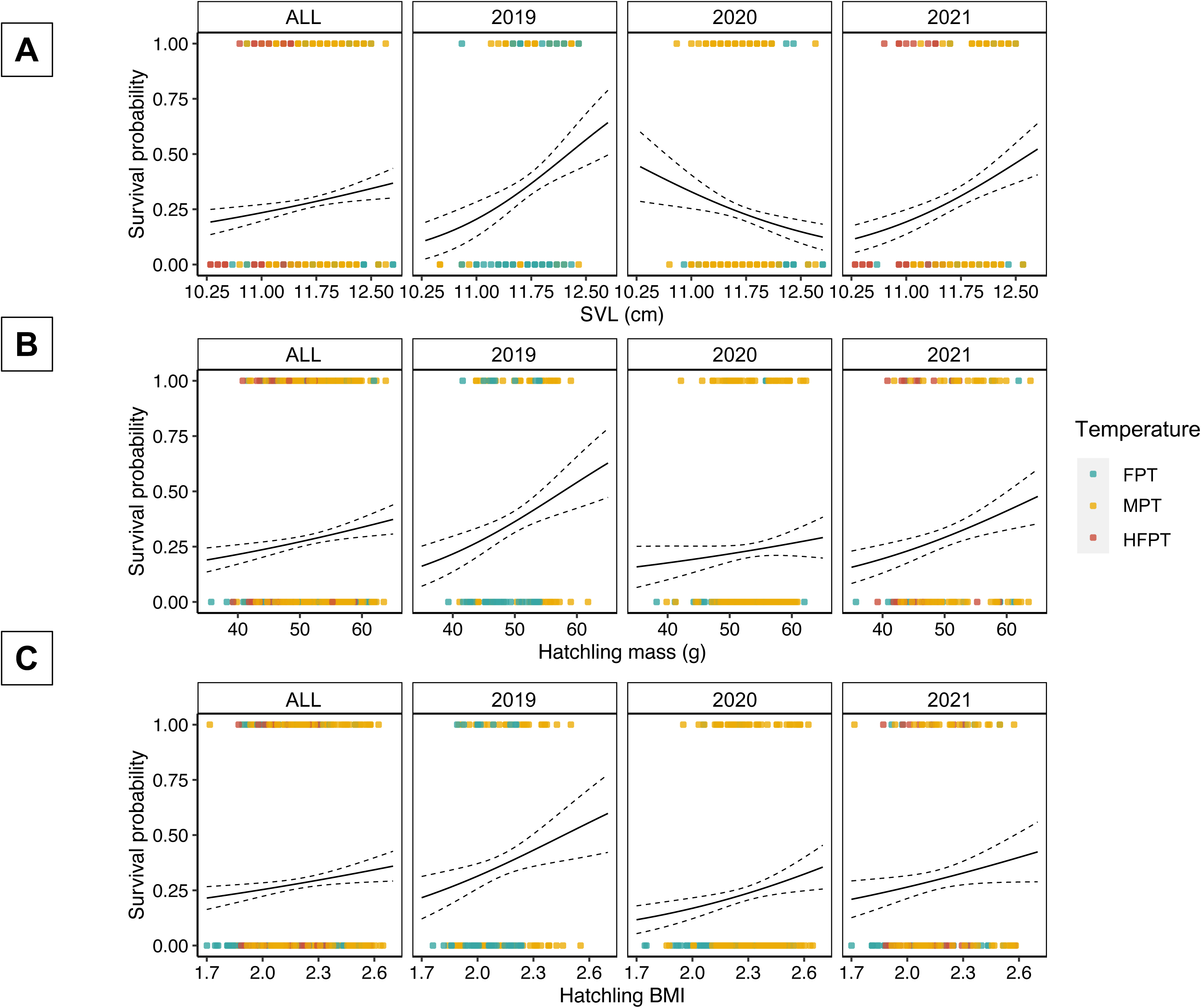
Associations between pre-winter survival status and temperature-related hatchling traits. Pre-winter survival status with respect to (A) hatchling snout-vent length (SVL), (B) body mass, and (C) body mass index (BMI). Solid line depicts the fit of a generalized linear model of pre-winter survival status with a binomial error distribution and each morphological trait. Dotted lines represent ± 1 SE. ‘ALL’ column includes hatchlings from all three cohorts pooled.

### Energetic cost of development is minimized at male promoting temperatures

Incubation temperature may alternatively exert lasting effects on hatchling survival via influences on physiological characteristics not necessarily reflected in morphology such as post-hatching energy reserves. Variation in metabolic rate was best explained by temperature and egg mass (ß_MPT_ = –0.0004 ± 0.0002, ß_eggmass_ = 0.0008 ± 0.0001, AICc = –339.48, *w_i_* = 0.64), though the temperature effect was not in the expected direction and the effect size was relatively small (**Figure 5A**). Incubation duration, on the other hand, was strongly associated with temperature (ß_MPT_ = –13.14 ± 0.33, AICc = 180.24, *w_i_* = 0.66), with individuals incubated at MPT hatching ∼13 days earlier than those incubated at FPT (**Figure 5B**). As a result, developmental cost, quantified as the product of embryonic metabolic rate and incubation duration (Marshall et al., 2020), was strongly influenced by temperature (ß_MPT_ = –3.01 ± 0.21, ß_eggmass_ = 0.09 ± 0.02, AICc = 130.07, *w_i_* = 0.74; **Figure 5C**). Embryos at FPT incurred a greater energetic cost of development, largely due to increases in incubation duration (**Figure 5D**).

**Figure 5.**
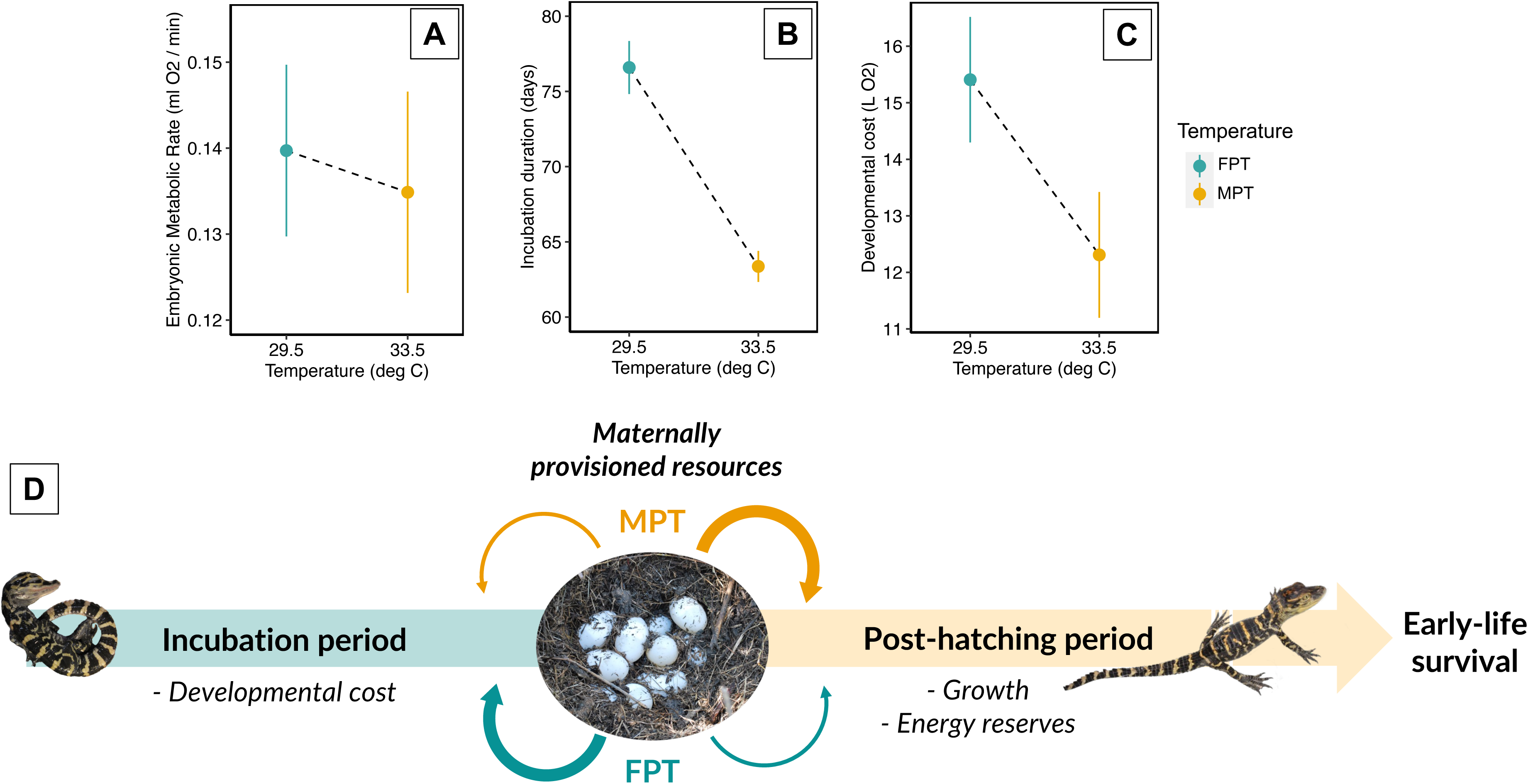
Temperature-dependent energetic cost of development as a potential mediator of variation in early life survival. (A) Embryonic metabolic rate (ml O_2_/min) measured at stage 26 under the same temperature conditions as those during incubation. (B) Incubation duration (days) measured as the time between estimated oviposition date and initiation of hatching. (C) Developmental cost measured as the product of embryonic metabolic rate and incubation duration (L O_2_ consumed during development if metabolic rate were constant). (D) Schematic depicting hypothesized relationship between incubation temperature, developmental energetic demands, and early-life survival.

## Discussion

Incubation temperature exerts a strong influence on the subsequent survival of hatchling alligators, with temperatures promoting male development conferring greater survival compared to those producing females. Importantly, the effects of incubation temperature persist in sex-reversed individuals, suggesting that temperature, rather than phenotypic sex drives variation in survival. When coupled with the observation that males reach age at first reproduction approximately a decade after females (Zajdel et al., 2019), our findings offer convincing empirical support for the hypothesis that differential survival-to-maturity underlies the adaptive value of TSD. Within this theoretical framework, incubation temperatures resulting in higher juvenile survivorship are predicted to produce the sex reaching maturity latest (Schwanz et al., 2016). Previous reports have shown that TSD species tend to display greater sex biases in age at maturity when compared to species with genetic sex determination (Bókony et al., 2019; Schwanz et al., 2016), yet experimental studies assessing incubation temperature effects on survivorship within this context are sparse.

It is thought that TSD evolved independently across different taxonomic groups and likely represents a convergent outcome stemming from different selective pressures (Janzen & Phillips, 2006; Sarre et al., 2011; Valenzuela & Lance, 2004). In contrast to the findings reported here in which incubation temperature affects survivorship, TSD in jacky dragons likely arose due to incubation temperature-driven variation in body size and consequent reproductive success (Warner & Shine, 2007, 2008). In contrast to many turtle and crocodilian species which display 10– to 20-fold longer lifespans, jacky dragons are among the shortest lived TSD reptiles and lack substantial sex biases in age at maturity (Warner & Shine, 2008). Given that most TSD reptiles are longer lived and display stark sex biases in age at maturity (Bókony et al., 2019), it is possible that incubation temperature affects juvenile survival more broadly and represents the predominate evolutionary explanation for the adaptive maintenance of TSD in reptiles. In support of this idea, common snapping turtle (*Chelydra serpentina*) hatchlings incubated at warmer, female-promoting temperatures were shown to have higher survivorship during their first year in experimental ponds when compared to hatchlings from lower, male-promoting temperatures (Janzen, 1995). In contrast to crocodilians, in most turtles, including *C. serpentina,* females typically reach maturity later than males and the increased survival of female hatchlings observed by Janzen is aligned with the survival-to-maturity hypotheses (Christiansen & Burken, 1979). Sexual size dimorphism, a trait related to differential age at maturity, has also been linked to mode of sex determination across reptiles lending further indirect support to the survival-to– maturity hypothesis (Katona et al., 2021). However, additional studies examining the extent to which incubation temperature influences juvenile survival and the directionality of this relationship with respect to sex biases in age at maturity are needed to generalize across reptiles exhibiting TSD.

Incubation temperature exerts clear effects on hatchling morphology, with incubation at male-promoting temperatures generally resulting in larger and more massive hatchlings when compared to both cooler and warmer temperatures. Consistent with our findings, a previous study demonstrated that incubation at male-promoting temperatures results in heavier hatchling alligators with more residual yolk stores, even after considering variation in egg size (Bock et al., 2021). Developmental cost theory predicts that the trade-off between temperature-mediated variation in incubation duration and metabolic rate is optimized at taxon-specific temperatures, at which conversion of maternal resources into offspring mass is most efficient (Marshall et al., 2020). Our findings indicate that incubation at male-promoting temperature minimizes developmental cost and likely contributes to observed variation in hatchling mass and residual yolk stores. Yet, the specific organismal traits that mediate the influence of incubation temperature on survivorship appear more ambiguous as relationships between hatchling morphology and survivorship varied across years. The effects of offspring size on survival are known to be context dependent in other reptiles, with size conferring survival advantages in some years and not others (Ferguson & Fox, 1984; Olsson & Madsen, 2001; Sinervo et al., 1992). In addition, incubation temperature is commonly reported to affect other hatchling traits (Noble et al., 2018) including hatchling behavior (Miller et al., 2020; Nichols et al., 2019), immune function (Leivesley & Rollinson, 2021; Treidel et al., 2016), and growth trajectories (Deeming & Ferguson, 1989; Marcó et al., 2010; Piña et al., 2007; Rhen & Lang, 1995), across diverse reptile species. It is intriguing to consider that the energetic cost of development may serve as a common underlying mechanism contributing to multiple fitness-related thermosensitive traits, but future studies directly linking developmental energetics to physiological traits and subsequent hatchling survival are required to test this hypothesis.

The present study does not address how long the influence of incubation temperature on survival persists beyond the first year of life. However, hatchlings from male-promoting temperatures displayed enhanced survival in both pre-winter and post-winter periods, suggesting that the effect is not limited to one season. Whereas long-term monitoring of individuals is required to determine the proportion of hatchlings reaching reproductive age, our analysis of reported sex ratios across different size classes suggests a marked shift from female– to male–biased sex ratios occurring during the hatchling-to-juvenile transition. The resulting male bias is then maintained through the juvenile-to-adult transition, resulting in broadly observed male skews in adult populations. A long-term mark recapture study in the same alligator population from which hatchlings in the 2020 and 2021 experiments originated suggest apparent survival rapidly increases in juveniles and small adults relative to hatchlings, but not in a way that differs by sex (Lawson et al., 2022). When taken together, available evidence suggests strong influences of incubation temperature on early-life fitness are likely sufficient to drive differences in survival to maturity, even if these temperature effects wane over time. Our findings provide empirical support for the hypothesis that differential survival to maturity contributes to the adaptive value of TSD in the American alligator. Future studies which follow the survival and reproductive outcomes of individuals into adulthood and which further examine the underlying mechanisms driving temperature-dependent early-life survival will be critical to unraveling the adaptive implications of temperature-dependent sex determination, a long-standing mystery in the field of evolutionary biology.

## Replication statement

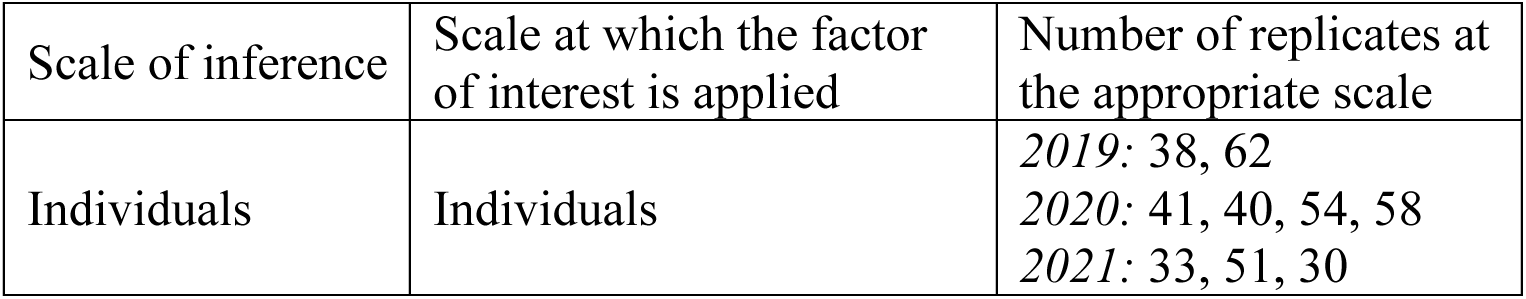

| Scale of inference | Scale at which the factor of interest is applied | Number of replicates at the appropriate scale |
| --- | --- | --- |
| Individuals | Individuals | 2019: 38, 62<br>2020: 41, 40, 54, 58<br>2021: 33, 51, 30 |

