## Supplemental information for "Differential early-life survival underlies the adaptive significance of temperature-dependent sex determination in a long-lived reptile"

**Supplemental Table 1.** American alligator sex ratios by size class.

| Reference | Site | Class | Size Range (TL) | Sex Ratio (% male) | Sample size (N) | Notes |
|---|---|---|---|---|---|---|
| Lance, Elsey, & Lang, 2000 | LA | J | 0.76-1.52m | 58 | 2936 | |
| Fuller, 1981 | NC | J | <1.8m | 66.7 | 25 | |
| | | A | >1.8m | 66.7 | 30 | |
| Bara, 1975 | SC | J | <0.9m | 57.3 | 482 | |
| | | J | 0.91-1.8m | 54.9 | 255 | |
| | | A | >1.8m | 60.8 | 51 | |
| Murphy, 1977 | SC | A | >1.8m | 78.9 | 90 | |
| Murphy & Coker 1983 | SC | A | 1.8-3.66m | 55.3 | 337 | |
| Brandt, 1991 | SC | A | 1.8-3.85m | 73 | 41 | |
| | | J | >1.8m | 78 | 145 | |
| Deitz, 1979 | FL | J | 0.41-1.08m | 61.3 | 727 | |
| Woodward et al. 1992 | FL | J | <0.61m | 66 | 1950 | |
| | | J | 0.61-1.21m | 60.7 | 1700 | |
| | | J | 1.22-1.82m | 54.3 | 300 | |
| | | A | 1.83-2.74m | 47.2 | 123 | |
| O'Neil, 1949 | LA | A | ~1.8m | 63.7 | 325 | |
| Chabreck, 1966 | LA | A | >1.8m | 60.4 | 234 | |
| Palmisano et al., 1973 | LA | J | 1.2-1.8m | 70.4 | 108 | |
| | | A | >1.8m | 83 | 195 | |
| Joanen, McNease, & Linscombe, 1974 | LA | J | 1.2-1.8m | 62.5 | 248 | |
| | | A | >1.8m | 67.9 | 595 | |
| Nichols & Chabreck, 1980 | LA | J | 0.6-1.8 | 59.8 | 1571 | |
| | | A | >1.8m | 68.4 | 57 | |
| Taylor, Kinler, & Linscombe, 1991 | LA | A | >1.8m | 56 | 5206 | |
| Rootes & Chabreck, 1992 | LA | J | 0.45-1.2m | 63.8 | 4151 | |
| Kinler & Taylor, 1992 | LA | J | <1.8m | 58 | 2337 | |
| | | A | >1.8m | 72 | 1836 | |
| Rhodes & Lang 1996 | SC | H | | 42.4 | 648 | Empirical |
| Elsey & Lang, 2014 | LA | H | | 28.1 | 6226 | Empirical |
| Ferguson & Joanen 1982 | LA | H | | 16.7 | 8000 | Empirical |
| Bock et al., 2020 | FL | H | | 40.6 | | Predicted |
| | SC | H | | 30.6 | | Predicted |

**Supplemental Table 2.** Generalized linear mixed effects models (GLMM) of pre-winter survival.

Abbreviations: $T_{inc}$ = incubation temperature.

| **Pre-winter survival** | | | | | | |
|---|---|---|---|---|---|---|
| **2019 cohort** | | | | | | |
| **model** | **AICc** | **Delta AICc** | **AICc weight** | **Cumulative weight** | **Log-likelihood** | **K** |
| ~ Tinc | 134.940 | 0.000 | 0.730 | 0.730 | -63.260 | 4 |
| Null | 136.930 | 1.990 | 0.270 | 1.000 | -65.340 | 3 |
| **2020 cohort** | | | | | | |
| **model** | **AICc** | **Delta AICc** | **AICc weight** | **Cumulative weight** | **Log-likelihood** | **K** |
| ~ Tinc | 179.999 | 0.000 | 0.467 | 0.467 | -85.893 | 4 |
| ~ Tinc + sex | 181.866 | 1.868 | 0.184 | 0.651 | -85.773 | 5 |
| ~ Tinc + treatment | 182.029 | 2.030 | 0.169 | 0.820 | -85.854 | 5 |
| ~ Tinc + sex + treatment | 183.850 | 3.852 | 0.068 | 0.888 | -85.699 | 6 |
| Null | 184.685 | 4.686 | 0.045 | 0.933 | -89.279 | 3 |
| ~ sex | 185.499 | 5.500 | 0.030 | 0.963 | -88.643 | 4 |
| ~ sex + treatment | 186.295 | 6.296 | 0.020 | 0.983 | -87.987 | 5 |
| ~ treatment | 186.607 | 6.608 | 0.017 | 1.000 | -89.197 | 4 |
| **2021 cohort** | | | | | | |
| **model** | **AICc** | **Delta AICc** | **AICc weight** | **Cumulative weight** | **Log-likelihood** | **K** |
| ~ Tinc | 141.223 | 0.000 | 0.556 | 0.556 | -65.334 | 5 |
| Null | 141.672 | 0.449 | 0.444 | 1.000 | -67.727 | 3 |

**Supplemental Table 3.** Generalized linear mixed effects models (GLMM) of post-winter survival. Abbreviations: $T_{inc}$ = incubation temperature.

| Post-winter survival | | | | | | |
|---|---|---|---|---|---|---|
| **2019 cohort** | | | | | | |
| **model** | **AICc** | **Delta AICc** | **AICc weight** | **Cumulative weight** | **Log-likelihood** | **K** |
| ~ Tinc | 75.630 | 0.000 | 0.850 | 0.850 | -33.610 | 4 |
| Null | 79.130 | 3.490 | 0.150 | 1.000 | -36.440 | 3 |
| **2020 cohort** | | | | | | |
| **model** | **AICc** | **Delta AICc** | **AICc weight** | **Cumulative weight** | **Log-likelihood** | **K** |
| ~ Tinc | 131.290 | 0.000 | 0.549 | 0.549 | -61.539 | 4 |
| ~ Tinc + sex | 133.281 | 1.991 | 0.203 | 0.752 | -61.480 | 5 |
| ~ Tinc + treatment | 133.389 | 2.099 | 0.192 | 0.944 | -61.534 | 5 |
| Null | 137.093 | 5.803 | 0.030 | 0.974 | -65.483 | 3 |
| ~ sex | 139.162 | 7.872 | 0.011 | 0.985 | -65.475 | 4 |
| ~ treatment | 139.174 | 7.884 | 0.011 | 0.996 | -65.481 | 4 |
| ~ sex + treatment | 141.017 | 9.727 | 0.004 | 1.000 | -65.348 | 5 |
| **2021 cohort** | | | | | | |
| **model** | **AICc** | **Delta AICc** | **AICc weight** | **Cumulative weight** | **Log-likelihood** | **K** |
| ~ Tinc | 62.569 | 0.000 | 0.669 | 0.669 | -26.007 | 5 |
| Null | 63.980 | 1.412 | 0.331 | 1.000 | -28.881 | 3 |

**Figure S1.**

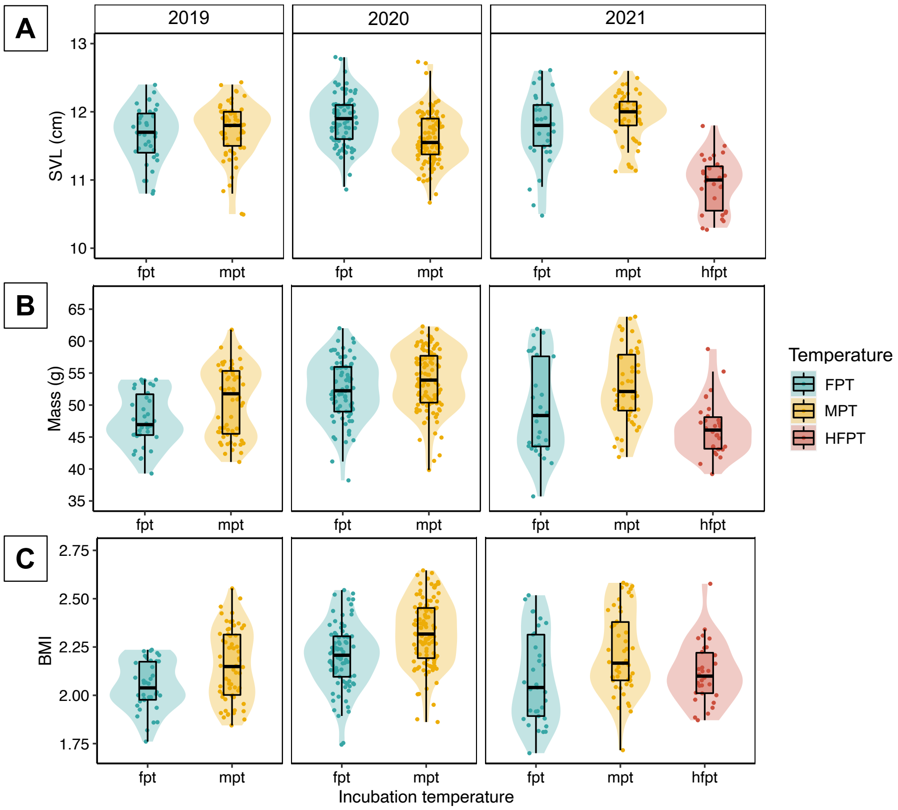


**Supplemental Figure 1.** Incubation temperature effects on hatchling morphometric traits – (A) snout-vent length (SVL), (B) body mass, (C) body mass index (BMI; mass/[2*SVL]) – measured shortly after hatching. In boxplots, central line indicates median, box indicates interquartile range (IQR), and vertical lines indicate maximum and minimum values or first and third quartile ± 1.5*IQR. Shaded violins depict kernel density of each group's trait value distribution. Abbreviations: FPT = female-promoting temperature, MPT = male-promoting temperature, HFPT = high female-promoting temperature.
